## Supplementary material for "Heron: A Knowledge Graph editor for intuitive implementation of python based experimental pipelines": Supplemnetary Material

**A**

```
Heron
    Heron
        Operations
        communication
            forwarders.py
            sink_com.py
            sink_worker.py
            socket_for_serialization.py
            source_com.py
            source_worker.py
            ssh_com.py
            ssh_info.json
            transform_com.py
            transform_worker.py
        docs
        gui
            create_new_node.py
            editor.py
            fdialog.py
            message.py
            node.py
            operations_list.py
            save_node_state.py
            settings.py
            ssh_info_editor.py
            visualisation_dpg.py

        resources
        templates
        settings_default.json
        constants.py
        general_utils.py
```

**C**

```
My_awesome_Nodes_repo
        README.md
        Sources
                Vision
                        __top__
                                ignore.gitignore
                        Weird_Camera
                                weird_camera_com.py
                                weird_camera_worker.com
        Transforms
                Motion
                        __top__
                                ignore.gitignore
                        Super_Motor_Controller
                                Maybe_Another_Folder
                                        another_script.py
                                super_motor_controller_com.py
                                super_motro_controller_worker.py
        Sinks
                Saving
                        __top__
                                ignore.gitignore
                        Saving_CSVs
                                some_other_script.py
                                saving_csvs_com.py
                                saving_csvs_worker.py
                General
                        __top__
                                ignore.gitignore
                        Some_Other_Sink_Node
                                some_other_sink_node_com.py
                                some_other_sink_node_worker.py
```

**B**

```
Base_repository_folder
    Sources / Transforms / Sinks
        Subcategory
            __top__
                ignore.gitignore
        Name_of_Node
            com_script.py
            worker_script.py
```

Supplementary Figure 1. Heron folder structures. A. The Heron folder structure together with the basic scripts that comprise Heron's core functionality. B. The Heron Node repository folder structure (to be found in the Operations folder for each group of Nodes in the same repository). C. Anexample of a repository with four Nodes (the Weird_Camera Source in subcategory Vision, the Super_Motion_Controller Transform in subcategory Motion and the Sinks Saving_CSVs in Saving and Some_Other_Sink_Node in General. The __top__ folder with the file ignore.ignore git file is required to allow Heron to create the list of Nodes presented to the users live as it's folder structure is updated with new Node folders.

```
import os
import sys
from os import path

current_dir = path.dirname(path.abspath(__file__))
while path.split(current_dir)[-1] != r'Heron':
    current_dir = path.dirname(current_dir)
sys.path.insert(0, path.dirname(current_dir))

from Heron import general_utils as gu
Exec = os.path.abspath(__file__)


BaseName = 'Awesome Transform Node'

NodeTooltip = 'Information about the Node’

NodeAttributeNames = ['Parameters',
                      'Something In',
                      'Something Out',
                      'Something Else Out Dict']

NodeAttributeType = ['Static', 'Input',
                      'Output', ‘Output’]

ParameterNames = ['Visualisation',
                  'Parameter 1', 'Parameter 2']

ParameterTypes = ['bool', 'str', 'list']

ParametersDefaultValues = [False,'True',
                           ['item 1', 'item 2']]
ParameterTooltips = [‘Info on Vis’, ‘Info on
                     Parameter1’, ‘Info on
                     Parameter2’]
InOutTooltips = [’Info on Something In’,
                 ‘Info on Something Out’,
                 ‘Info on Something Else Out Dict’]


WorkerDefaultExecutable = \
os.path.join(os.path.dirname(Exec),
             'transform_template_worker.py')

if __name__ == "__main__":
    transform_template_com = \
gu.start_the_transform_communications_process\
(NodeAttributeType, NodeAttributeNames)
    gu.register_exit_signals(\
     transform_template_com.on_kill)
    transform_template_com.start_ioloop()
```

Code allowing Heron to find the script without package installation. This should not be modified.

Here users enter the values (blue) of the variables that define the Node’s Name, Inputs, Outputs and Parameters. Heron will automatically generate the Node’s GUI on Heron’s Node Editor using these values.

Code that deals with the starting of the com process. This should not be modified.

Box 1A. An abbreviated and annotated template for the com script of a Transport Operation

```python
import sys
from os import path
import numpy as np

current_dir = path.dirname(path.abspath(__file__))
while path.split(current_dir)[-1] != r'Heron':
    current_dir = path.dirname(current_dir)
sys.path.insert(0, path.dirname(current_dir))

from Heron.communication.socket_for_serialization import Socket
from Heron import general_utils as gu, constants as ct
from Heron.gui.visualisation import VisualisationDPG


need_parameters = True
global_var_1: str
global_var_2: str

vis: VisualisationDPG


def initialise(worker_object):
    global vis
    global need_parameters
    global global_var_1
    global global_var_2


    try:
        parameters = worker_object.parameters
        global_var_1 = parameters[1]
        global_var_2 = parameters[2]
    except:
        return False


    visualisation_type = 'Value'
    buffer = 20
    vis = VisualisationDPG(_node_name=worker_object.node_name,
                           _node_index=worker_object.node_index,
                           _visualisation_type=visualisation_type,
                           _buffer=buffer)

     worker_object.savenodestate_create_parameters_df(
          parameter_var_1=global_var_1,
          parameter_var_2=global_var_2,
          parameter_var_3=global_var_3,
          parameter_var_4=global_var_4)

    return True


def work_function(data, parameters):
    global vis
    global global_var_1
    global global_var_2

    try:
        global_var_1 = parameters[1]

        vis.visualisation_on = parameters[0]
    except:
        pass

    topic = data[0]
    print(topic)

    message = data[1:]
    data = Socket.reconstruct_array_from_bytes_message(message)

    # Now do stuff

    savenodestate_update_substate_df(image__shape=message.shape)

    vis.visualise(message)

    result = [message, np.array([ct.IGNORE])]

    return result

def on_end_of_life():
    global vis

    vis.kill()


if __name__ == "__main__":
    worker_object = \
gu.start_the_transform_worker_process(\
     work_function=work_function,
     end_of_life_function=on_end_of_life,
     initialisation_function=initialise)

    worker_object.start_ioloop()
```

Code allowing Heron to find the script without package installation. This should not be modified.

Global variables is just one of the possible ways for the intialise, work_function and on_end_of_life functions to keep state over multiple calls.

The initialise function is called at the start of the process recurently until it returns True. The worker_function will not be called until the initialise function returns True. Here goes the code that deals with the initialisation of the worker process (like grabbing the values of the parameters from the Node's GUI).

The worker_function is where the main code of the Operation should be placed. It will be called by Heron every time there is data coming into the Node. It will return a list of numpy arrays. The items of that list is what the Node sends from its outputs (each item corresponding to one output).

The end_of_life function will be called by Heron when the process is about to terminate so any code to clean up goes here.

This is the code that is called by the com process to start the worker process. This should not be modified.

Box 1B. An abbreviated and annotated template for the worker script of a Transport Operation

### Worker_function Code

#### Trial Generator

```
def work_function(data, parameters):
    global vis
    global lengths_block
    global current_block
    global reward_contingencies
    global correct_licks

    vis = parameters[0]

    message = data[1:]   # data[0] is the topic
    message = Socket.reconstruct_array_from_bytes_message_cv2correction(message)

    # If the message comes from the Trial controller
    if len(message) == 2:
        # The Trial controller sends [previous_stim, correct_port_licks]
        previous_stim = message[0]
        correct_port_licks = message[1]

        # Add the number of correct licks to the current running sum
         correct_licks[previous_stim]  =  correct_licks[previous_stim]  +
                                          correct_port_licks

        # If the number of correct licks reaches the block length of that stim then
        # swap block and zero the running sum of licks for that stim
        if correct_licks[previous_stim] ==
          lengths_block[current_block[previous_stim]][previous_stim]:
            create_new_block_sizes(previous_stim)
            temp = copy.copy(current_block[previous_stim])
            current_block[previous_stim] = int(not current_block[previous_stim])
            correct_licks[previous_stim] = 0
           if vis:
                 print('Changing block of stim {}, from block {} to block
                   {}'.format(previous_stim, temp, current_block[previous_stim]))
                 print('Current block lengths = {}'.format(lengths_block))

    current_stim = np.random.randint(0, 4)
    current_correct_reward_port_probability =
             reward_contingencies[current_block[current_stim]][current_stim]
    current_correct_reward_port = np.random.binomial(1,
                                       current_correct_reward_port_probability)

    result = [np.array([current_stim,
              current_correct_reward_port,
              current_block[current_stim]])]

    return result
```

Box 2A. The work function of the Trial Generator Operation

#### Trial Controller

```
def work_function(data, parameters):
    global vis
    global respond_after_lick
    global start_delay
    global odour_window
    global pre_response_delay
    global response_window
    global reward_window
    global inter_trial_window
    global trial_number
    trial_number += 1

    try:
        vis = parameters[0]
    except:
        pass
```

```
    # Get message in from previous Node
    message = data[1:]   # data[0] is the topic
    message = Socket.reconstruct_array_from_bytes_message_cv2correction(message)
    stim = message[0]
    reward_port = message[1]
    block_of_stim = message[2]

    if vis:
        print('============= Starting Trial {} ============'.format(trial_number))
        print('Current Stim = {}, current correct reward port = {}'.format(stim,
                                                                          reward_port))

    start_trial_time = now()
    if vis:
        print(' ooo Waiting Start Delay {}'.format(start_delay))
    gu.accurate_delay(start_delay * 1000)

    open_stim_time = now()

    if vis:
        print(' ==== Arduino Open Stim {}'.format(stim))
    arduino_serial.write(stim_on_commands[stim])

    if vis:
        print('ooo Starting Odour Delay {}'.format(odour_window))
    gu.accurate_delay(odour_window * 1000)

    close_stim_time = now()
    if vis:
        print(' === Arduino Close Stim {}'.format(stim))
    arduino_serial.write(stim_off_commands[stim])

    if vis:
        print('ooo Starting Pre-Response Delay {}'.format(pre_response_delay))
    gu.accurate_delay(pre_response_delay * 1000)

    correct_port_lick, start_response_time = read_arduino(reward_port)

    if respond_after_lick and correct_port_lick:
        start_reward_time, end_reward_time = reward(correct_port_lick, reward_port)
    elif respond_after_lick and not correct_port_lick:
        if vis:
            print(" ooo Correct port hasn't been licked = No reward")
        start_reward_time = datetime.datetime.now()
        end_reward_time = start_reward_time
    elif not respond_after_lick:
        start_reward_time, end_reward_time= reward(correct_port_lick, reward_port)

    if vis:
        print('ooo Starting Response Delay {}'.format(reward_window))
    gu.accurate_delay(reward_window * 1000)
    iti = np.random.uniform(inter_trial_window[0], inter_trial_window[1])

    if vis:
        print('ooo Starting ITI Delay {}'.format(iti))
    gu.accurate_delay(iti * 1000)
    end_iti_time = now()

    result = [np.array([stim, correct_port_lick]),
              np.array([start_trial_time, open_stim_time, close_stim_time,
                        start_response_time, start_reward_time, end_reward_time,
                        end_iti_time, str(stim), str(reward_port),
                        str(block_of_stim), str(correct_port_lick)])]

    if vis:
        print(' ooo Stim and Correct Port Lick = {}'.format(result[0]))
        print('============== Ended Trial {}================'.format(trial_number))
        print('----------------------------------------------')

    return result
```

Box 2B. The worker function of the Trial Controller Operation

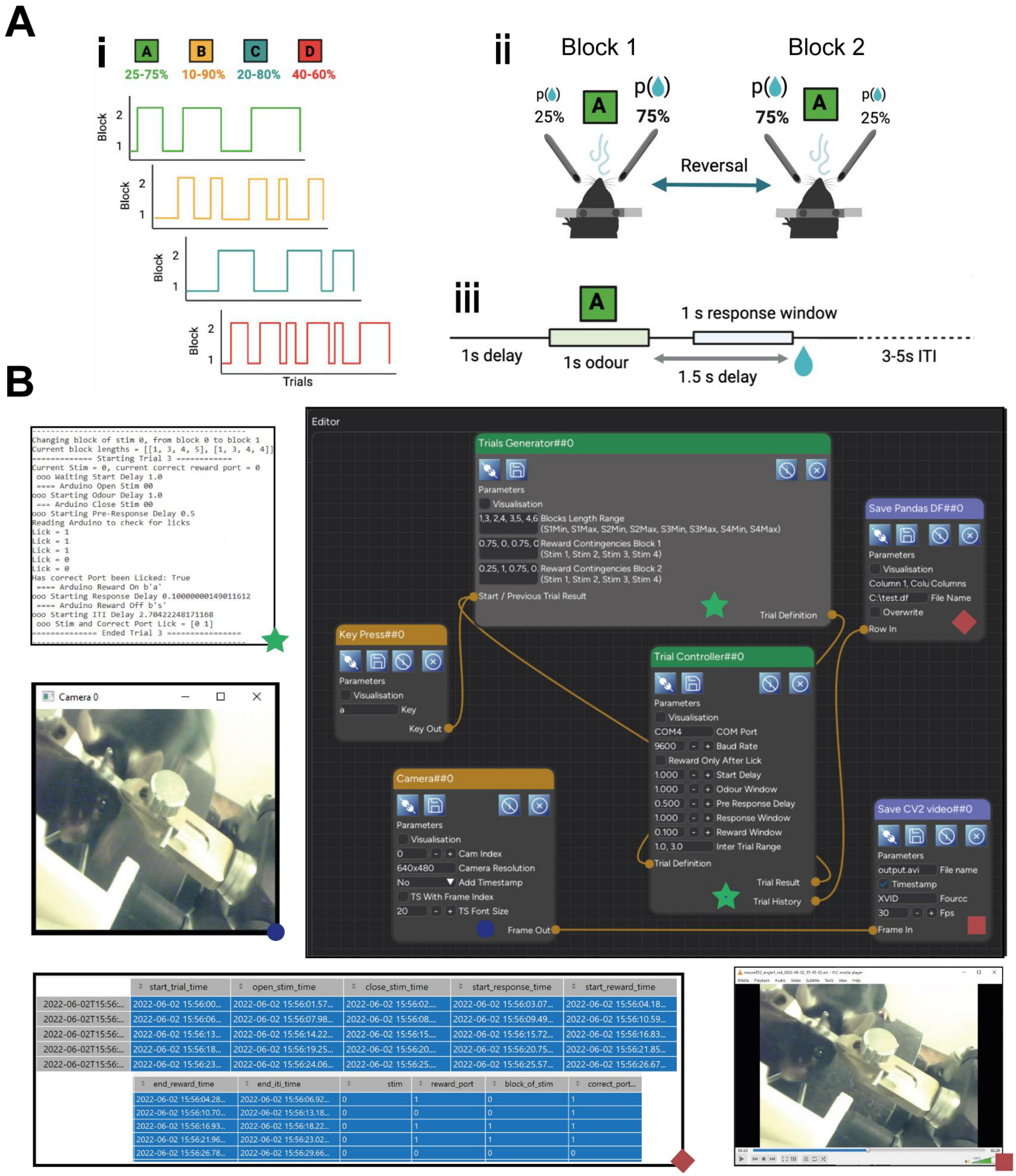


Supplementary Figure 2. The Probabilistic Reversal Learning experiment. A. A schematic of the task. i) The block structure for the 4 odours (stimuli) showing the transitions over trials between block 1 and block 2 for each odour. These can also be seen (and set for each session) in the 'Reward Contingency Block 1' and 'Reward Contingency Block 2' parameters of the 'Trials Generator' Node (see B). ii) A diagrammatic description of what a block transition for one odour (in this case, odour A) means. iii) The dynamics of a single trial. All the timings can be seen (and set) in the parameters of the 'Trial Controller' Node. B. A snapshot of the experiment running. Top right image is Heron showing the full Knowledge Graph. Images to the left of the Heron GUI are the outputs of the different Node visualisations showing live while Heron is running. The images below the GUI are snapshots of the saved results for the two Sinks ('Save Pandas DF' and 'Save FFMPEG Video'). Which output is generated by which Node is marked with green stars, blue circles and red squares and diamonds at the bottom of the Node and the corresponding image.

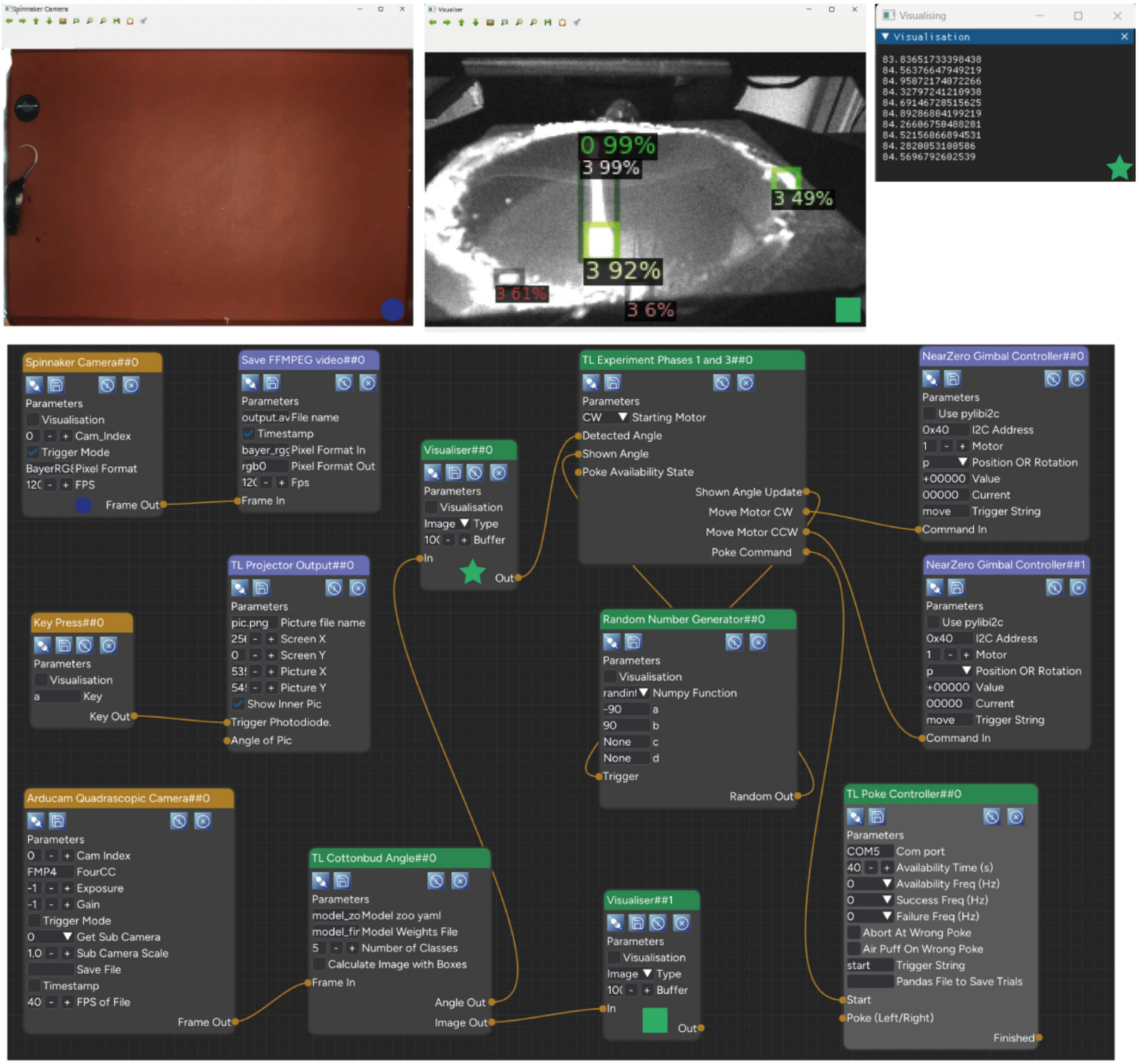


Supplementary Figure 3. Fetching the cotton bud experiment as controlled by Heron. The bottom image is the Heron Knowledge Graph. The top three images are the visualisations of Nodes “Spinnaker Camera”, “Visualiser##0” and “Visualiser##1”. Which visualisation belongs to which Node is marked by a green star, a blue circle and a green square at the bottom of the Nodes and its corresponding visualisation image. Visualiser##0 is receiving its values from the Angle Out output of the “TL Cottonbud Angle” Node while “Visualiser##1” is receiving its frames from the Image Out output of the “TL Cottonbud Angle” Node. The “TL Cottonbud Angle” Node is receiving frames of the bottom of the cotton bud receptacle from the Frame Out output of the “Arducam Quadrascope Camera” Node and is using one of the algorithms in the Detectron2 package (which one is defined in the ‘Model zoo yaml’ parameter of the “TL Cottonbud Angle” Node ) to both calculate the angle of the cotton bud and to superimpose on the receiving frame the detected boxes of the cotton bud’s tips and of the cotton bud itself. The frame of the cotton bud is captured on a Jetson Nano computer, is passed onto the Windows machine where the Heron GUI is running, to be then passed to the WSL virtual machine inside the Windows machine for the deep learning algorithm to use it as input. The output of the algorithm (the angle and the boxes superimposed image) is then passed back to the Windows machine to be used by the logic of the experiment running in the “TL Experiment Phases 1 and 3” Node.

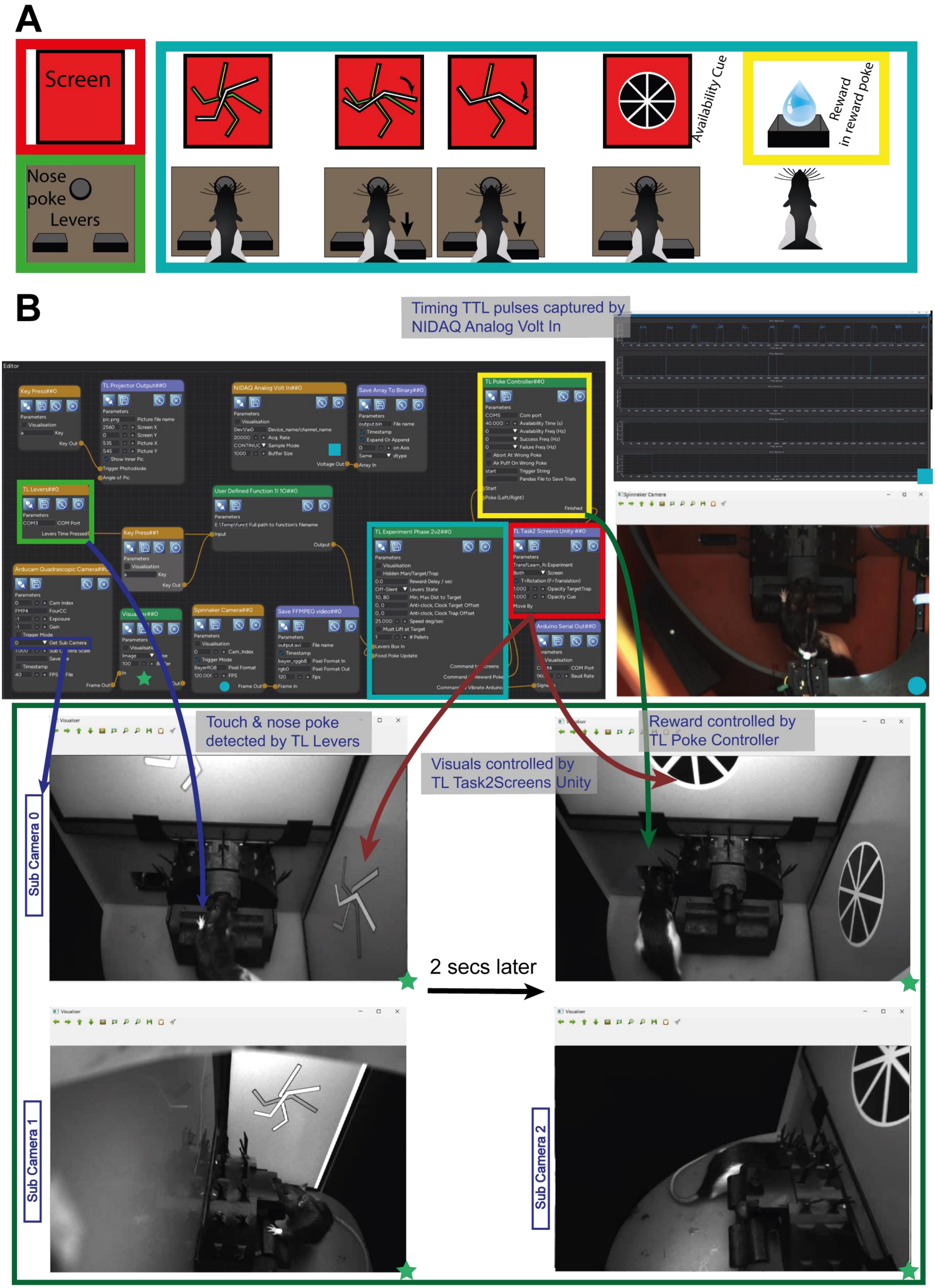
A
Screen
Nose poke
Levers
Availability Cue
Reward in reward poke
B
Timing TTL pulses captured by NIDAQ Analog Volt In
Touch & nose poke detected by TL Levers
Visuals controlled by TL Task2Screens Unity
Reward controlled by TL Poke Controller
Sub Camera 0
Sub Camera 1
Sub Camera 2
2 secs later
Editor
Key Press##0
TL Projector Output##0
NIDAQ Analog Volt In##0
Save Array To Binary##0
TL Poke Controller##0
TL Levers##0
User Defined Function 1I 1O##0
Key Press##1
TL Experiment Phase 2v2##0
TL Task2 Screens Unity ##0
Arducam Quadrascopic Camera##0
Spinnaker Camera##0
Save FFMPEG video##0
Arduino Serial Out##0
Spinnaker Camera
Visualiser

Supplementary Figure 4. Heron running the 'computer game for rats' experiment. A. The experimental design. The animal can access a nose poke with two levers box (left bottom, Light Green box) and can see two screens (represented here as one rectangle, left top, Red box). The experiment detects the animal's nose poke and presents three lines (white, yellow and green) on a red background. If the animal presses any of the two levers the white line rotates either clock or counter clock wise. If the white line is rotated to reach the green line then a reward availability cue (black spoked wheel) is presented and animated indicating to the animal that reward is available in a separate reward port. The detection of the nose poke and the levers is controlled by the "TL Levers##0" Node (the joystick of the game). The timing of the availability period, the detection of the animal at the reward port and the dispensing of reward is controlled by the "TL Poke Controller##0" Node. The logic of the game is controlled by the "TL Experiment Phase 2v2##0" Node which receives the input from the 'joystick' and decides what to show the animal and whether to reward it. Finally, the visuals of the game are controlled by the "TL Task2 Screens Unity##0" Node which receives commands from the logic Node and passes them to a Unity executable which deals with the proper presentation of the sprites and their animations. B. Heron with its live visualisations of the FLIR camera, the TTL pulses used for timing of the different events (as captured by a NIDAQ board (National Instruments)) and the different "sub cameras" parts of the Arducam 4 camera system. The four coloured boxed surrounding the Nodes "TL Levers##0" (Light Green), "TL Experiment Phase 2v2##0" (Light Blue), "TL Poke Controller##0" (Yellow) and "TL Task2 Screens Unity##0" (Red) are not shown on the Heron GUI but are used here to indicate which Node controls which part of the experimental design shown in A. The arrows show which Node is responsible for which aspect of the experiment as it appears on the captured video streams. The correspondence between the Nodes and their visualisation is marked with green stars, a blue circle and a blue square at the bottom of the Nodes and their corresponding visualisation windows. The green window shows two time points as seen by two of the three sub cameras used in the system. In the first time point (left column of bottom green rectangle, showing sub cameras 4 and 2) the animals is pressing the left lever making the white jagged line on the screens go towards the darker line (green on the coloured monitors). A couple of seconds later the animals has rotated the white line to touch the green one and the reward availability cue (dark spoked disk) is animating on the screens. The animal then rushes to receive its reward from the reward tray as shown on the second column of the sub camera captures.
